# Nanopore electro-osmotic trap for the label-free study of single proteins and their conformations

**DOI:** 10.1101/2021.03.09.434634

**Authors:** Sonja Schmid, Pierre Stömmer, Hendrik Dietz, Cees Dekker

**Affiliations:** Department of Bionanoscience, Kavli Institute of Nanoscience, Delft University of Technology, Van der Maasweg 9, 2629 HZ Delft, The Netherlands; NanoDynamicsLab, Laboratory of Biophysics, Wageningen University, Stippeneng 4, 6708 WE Wageningen, The Netherlands; Physik Department, Technische Universität München, Garching near Munich, Germany

## Abstract

Many strategies have been pursued to trap and monitor single proteins over time in order to detect the molecular mechanisms of these essential nanomachines. Single protein sensing with nanopores is particularly attractive because it allows label-free high-bandwidth detection based on ion currents. Here we present the Nanopore Electro-Osmotic trap (NEOtrap) that allows trapping and observing single proteins for hours with sub-millisecond time resolution. The NEOtrap is formed by docking a DNA-origami sphere onto a passivated solid-state nanopore, which seals off a nanocavity of a user-defined size and creates an electro-osmotic flow that traps nearby particles irrespective of their charge. We demonstrate the NEOtrap’s ability to sensitively distinguish proteins based on size and shape, and discriminate nucleotide-dependent protein conformations, as exemplified by the chaperone protein Hsp90. Given the experimental simplicity and capacity for label-free single-protein detection over the broad bio-relevant time range, the NEOtrap opens new avenues to study the molecular kinetics underlying protein function.

---

All vital functions in our cells are performed by proteins. Understanding protein function ultimately means understanding how these complex molecules behave over time, i.e., how they adopt multiple energetically accessible conformations in solution in a thermal heat bath, often driven by additional external energy sources such as ATP, light, ion gradients, etc. Much progress has recently been made in obtaining high-resolution 3D structures from cryo-electron microscopy, which is greatly informative but provides static snapshots, and hence the kinetics underlying protein function remain inaccessible. Single-molecule techniques can powerfully resolve molecular details which in bulk are masked by ensemble averaging, but they also face limitations such as a narrow temporal bandwidth (e.g. in Förster resonance energy transfer; FRET), interference from surface interactions (in atomic force microscopy), or the need for site-specific labeling (FRET dyes, handles for optical or magnetic tweezers). Scientists have developed new types of single-molecule traps aiming to overcome such shortcomings. For example, the anti-Brownian electro-kinetic (ABEL) trap^1,2^ can hold a dye-labeled protein for seconds using real-time electro-kinetic feedback control. Electrostatic fluidic traps were shown to trap unfolded dye-labeled proteins under very low salt conditions, and to detect their charge^3^. Plasmonic nanoapertures can trap unlabeled single proteins^4,5^, but control and reproducible fabrication is challenging. Dielectrophoretic long-term trapping was demonstrated with silica beads but not on the level of individual protein molecules^6^ while only recently first short-duration (≤ms) protein detection was shown^7^. Most of these methods require labeling of the proteins and sophisticated instrumentation, which limits their use to a handful of expert labs.

Here we present the Nanopore Electro-osmotic trap (NEOtrap), a nanopore-based device that is able to trap single unmodified proteins for up to hours, while recording conformation-sensitive currents at sub-millisecond resolution. We start by introducing the trapping concept that is based on docking a DNA-origami nanosphere onto a solid-state nanopore and subsequent trapping of single proteins. Next, we demonstrate the single-molecule sensitivity and high resolution of the NEOtrap by discriminating a variety of proteins based on their mass and shape. Data for various pore sizes, voltages, and ionic strengths inform on the trapping mechanism. Finally, we address protein function and present data that, remarkably, resolve nucleotide-dependent shifts in the conformations of the chaperone protein Hsp90, notably detected label-free and at the single-molecule level using the NEOtrap.

## Building the nanopore electro-osmotic trap

Single-protein trapping using the NEOtrap occurs in two steps (Fig. 1), (i) *in situ* assembly of the molecular trap and (ii) subsequent trapping of the protein of interest. The sensor consists of a solid-state nanopore, typically with a diameter d_pore_ ≈ 20 nm which is TEM-drilled into a 20 nm thick silicon-nitride membrane^8^ and subsequently coated with a ∼5 nm thin lipid bilayer for passivation^9^, thus resulting in a nanopore with a diameter d_coated pore_ = d_pore_ – 10nm ≈ 10nm (Fig. 1a). Voltage application (typically +100 mV) across the pore establishes a baseline ion current. The second ingredient of the NEOtrap is a charged and permeable nanostructure, in this case a 35 nm diameter DNA-origami sphere^10^. Once such a negatively charged sphere diffuses close to the nanopore, the electric field drives it to the pore electrophoretically, where it gets docked onto the pore entrance^11–13^ thus closing off a nanocavity. This leads to a characteristic and reproducible small current drop (∼20%), since origami nanostructures are permeable to ions^11,14^. The negative charge of the DNA phosphate backbone is screened by positive counter ions, and in the applied electric field, these counter ions migrate towards the cathode (upwards in Fig. 1a), which results in a strong hydrodynamic flux towards the pore. This nanofluidic effect is called electro-osmosis and is well studied in the context of surface charge in nanochannels and nanopores^15–18^ (see also Supplementary Notes 3.1). Here, however, the electro-osmosis is not surface driven – as the lipid bilayer surface is net neutral – but induced site-specifically by the DNA origami structure that acts as a nanoporous negatively charged sponge. The electro-osmotic flow (EOF) drives proteins with the surrounding solvent towards the nanopore. As a result, a single protein can be trapped in the nanocavity formed by the coated nanopore that is capped by the docked origami sphere.

**Figure 1.**
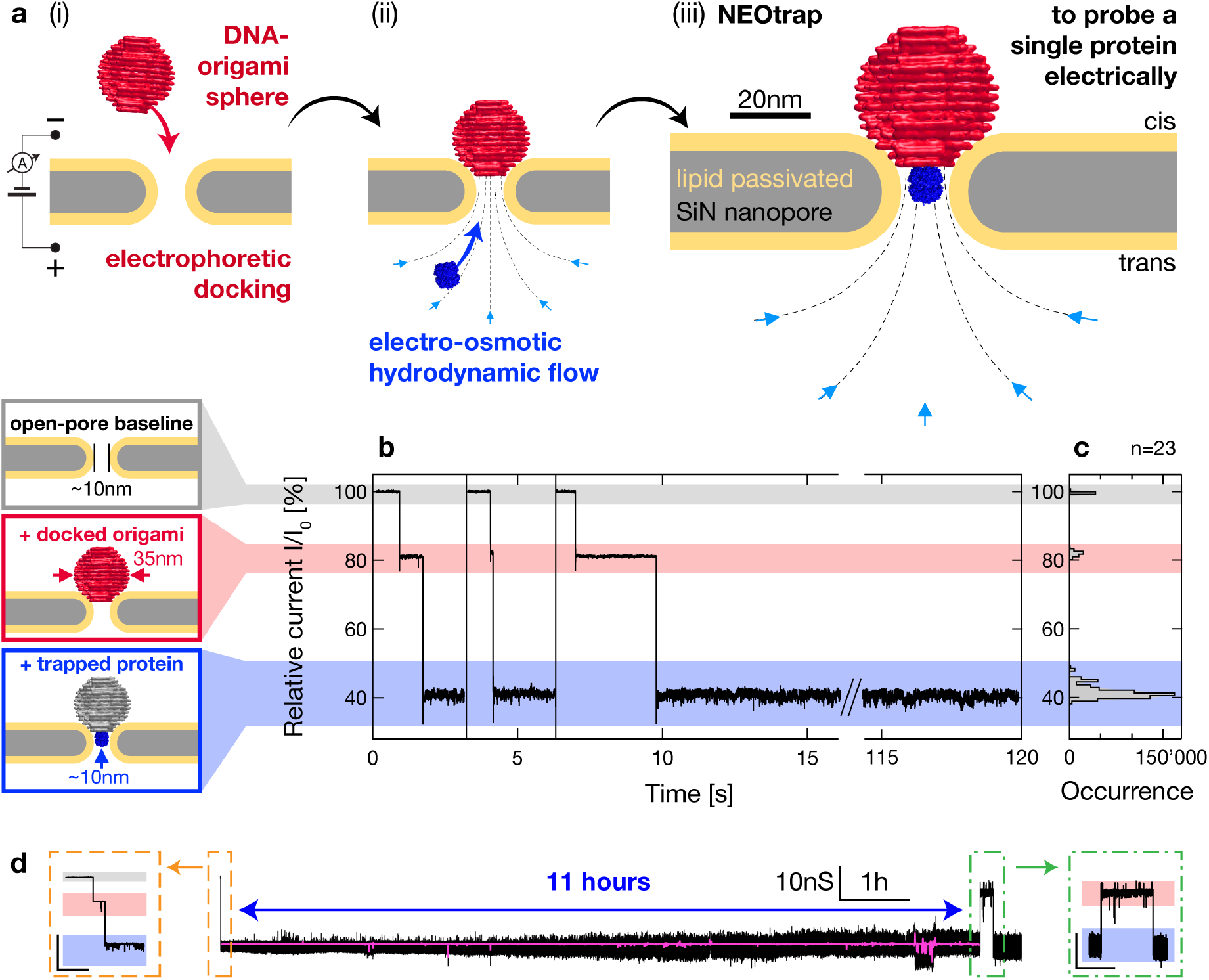
Working principle of the Nanopore Electro-Osmotic trap. **a** Illustration of the two-step trapping process: a DNA-origami sphere is electrophoretically docked onto a passivated solid-state nanopore, inducing hydrodynamic flow caused by electro-osmosis that facilitates protein trapping at the centre of the nanopore. ‘Cis’ and ‘trans’ reservoirs are defined in panel (iii). **b** Current recordings start with a low-noise open-pore baseline (gray color code); electrophoretic origami docking blocks ∼20% of the current (red); the induced electro-osmotic flow traps a ClpP protein leading to further current blockage (blue). Three consecutive origami docking events followed by protein trapping (at 100 mV) are shown. In between these events, the trapped proteins were manually released by voltage inversion (−100 mV). The displayed relative current is normalized by I_0_, the open-pore current (in the absence of a docked origami or a trapped protein) and 1 kHz low-pass filtered. More current traces are shown in Supplementary Figure 1. **c** Histogram of 23 current traces such as those in (b). **d** One ClpP protein was trapped and recorded for 11 hours. Insets show zoom views of ClpP trapping (orange, scale bar 10 nS vs. 10 s) and spontaneous escape after 11h (green, scale bar 10 nS vs. 500 s); gray, red, blue shading as in (b). The daylong recording was corrected for baseline drift and conductance increase over time (black: 1 kHz; pink: 100 Hz median). The unprocessed data are shown in Supplementary Fig. 2. The origami sphere in panels (a, b) was adapted from Ref.10.

The trapping process can be monitored in real time (Fig. 1b): current traces show transitions from the open pore current to a first partial blockade (due to the DNA sphere docking), and then to a second blockade associated with the trapping of a single protein into the cavity. This single-protein trapping is highly reproducible and comes with a characteristic current blockade, as shown in Fig. 1c and Supplementary Fig. 1. A single protein can be trapped reversibly for very long times. We demonstrated single-protein trapping up to 11 hours, see Fig. 1d and Supplementary Fig. 2. The NEOtrap is easy to assemble and can be reset simply by inverting the voltage, which releases the DNA origami sphere and cleans up the trap, thus preparing it for a next round of assembly and trapping. Indeed, we routinely use a single nanopore for hundreds of single-protein trapping events in succession. Furthermore, the EOF caused by the DNA-origami sphere allows to trap proteins irrespective of their charge (Supplementary Table 1). The data shown in Fig. 1, for example, were obtained for the negatively charged protein ClpP (net charge -55 q_e_), which is drawn into the NEOtrap *against* the electrophoretic driving force acting on it, proving that the EOF dominates, and indicating that the NEOtrap can be used for a variety of charged biomolecules. Overall, the data show that the NEOtrap allows for easy and versatile long-term trapping of single proteins, which provides ample observation time of unmodified proteins.

## Distinguishing single proteins by size and shape

In Fig. 2, we present the ability of the NEOtrap to distinguish individual proteins based on their current blockade signals observed in one and the same nanopore with a diameter d_pore_ = 20 nm. We compare proteins of different mass (from 50 to 325 kDa), size (4.5 to 13 nm), and shape, viz., avidin, Hsp90, ClpX, and ClpP. All trapped proteins yielded characteristic signals, with a well-defined single peak in the blockade current histogram of most proteins (Fig. 2b) – except for ClpX, which produces two peaks, as discussed below. As expected from volume-exclusion considerations, larger proteins produced deeper current blockades, and a linear dependence was observed between the main blockade peak and the molecular weight of the proteins (Fig. 2c). An approximately linear dependence can be expected for globular proteins, as larger-mass proteins occupy a larger volume in the electrolyte-filled pore, thus blocking the ion flow, accordingly. Vice versa, the relationship can also be used to estimate the mass of a trapped protein from the current blockade. While the capture rate was found to vary little in all cases, the trapped time (i.e., inverse escape rate) increased exponentially as a function of molecular weight (see Supplementary Fig. 3), consistent with theoretical predictions (Supplementary Notes 3.2 and 3.3).

**Figure 2.**
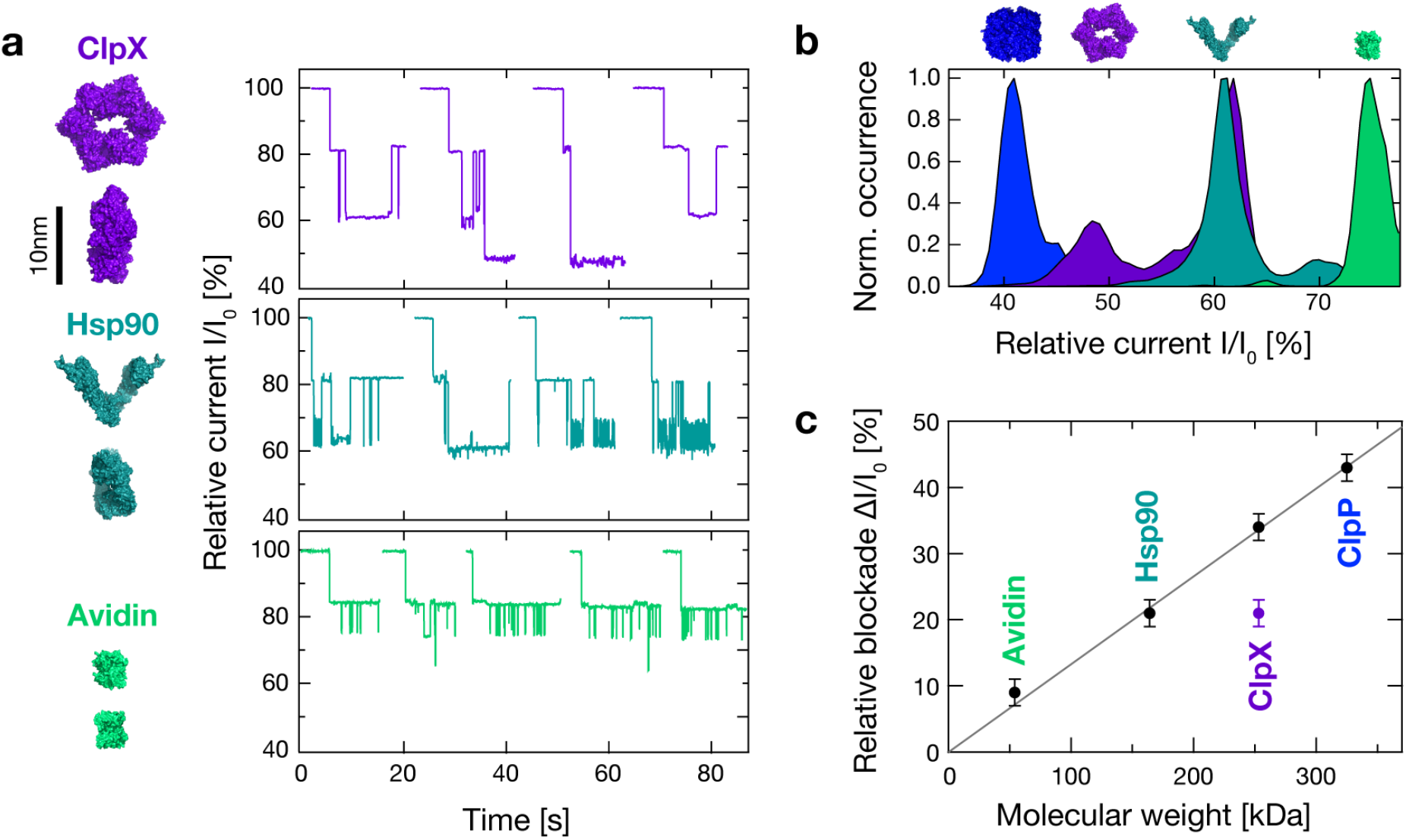
Mass- and shape-dependent single-protein identification with the NEOtrap. **a** Representative current recordings showing protein trapping events of ClpX, Hsp90, avidin for a 20 nm pore at 100 mV, with corresponding structures in front- and side-view (left images). The current is normalized by I_0_, the open-pore current in the absence of a docked origami or a trapped protein and 1 kHz low-pass filtered. **b** Relative current histogram of 10-30 origami docking events with trapped ClpP, ClpX, Hsp90, avidin, colour-coded as in (c). **c** Relative current blockade ΔI/I_0_ of each protein as a function of molecular weight. The line represents a linear fit through the black markers (slope = 0.13±0.01 %/kDa; y-axis intercept = 1.5±2.2 %. The experiment was repeated with two pores in three separate experiments showing a similar linear mass-dependence. For ClpX, two data points are observed, due to its disk shape, as discussed in the text.

Interestingly, the NEOtrap does not merely measure protein mass, but can also be used to sense protein shape. This can be seen from the data on disk-shaped ClpX protein, where the current histogram (Fig. 2b) shows not one but two peaks, ΔI/I_0_ = 34±2% and 21±2%, with the second peak at a much smaller blockade (i.e., higher current level). We attribute the two current levels to two different orientations of the ClpX structure – parallel and perpendicular to the ion flow – in line with previous nanopore observations of orientation-specific ion-current signals by Yusko *et al*^19^. Summing up, the data in Fig. 2 demonstrates the NEOtrap’s potential to identify proteins based on their mass, size, and shape.

## Trapping dependence on pore size, voltage, and ionic strength

The pore size has a strong effect on the observed single-protein trapping behaviour, as directly visible from the ClpP traces in Fig. 3a. Three regimes can be distinguished, viz., small (S), intermediate (M), and large (L) sizes of the cavity after lipid-bilayer coating of the nanopores. A cavity with d_coated pore_ ≤ 10 nm (S) is too small to accommodate ClpP and many short-duration events are observed. These are attributed to ClpP proteins that bump onto the bottom rim of the cavity (in Fig. 3a), whereupon they return to the same reservoir. Note also that the magnitude of electro-osmotic flow is weakest for these small pore diameters (see Supplementary Notes 3.2). For cavity diameters of d_coated pore_ = 10-15 nm (M), one ClpP can get stably trapped for hours (Fig. 1d). Since ClpP’s size fits tightly into the cavity, the electro-osmotic flow is substantially blocked upon trapping, and no additional (bumping) events are observed. The presence of a protein thus limits the trapping strength for capturing additional proteins, providing a mechanism for self-regulation of *single*-protein trapping in this regime. By contrast, for cavity sizes of d_coated pore_ > 16 nm (L), multiple ClpP proteins are trapped consecutively, leading to a staircase behaviour in the current traces (Fig. 3a bottom). Here, each individual ClpP protein leads to the same conductance blockade of 7.1±1.0 nS, for up to three ClpP’s that fit completely into the nanocavity, while the blockade for the forth trapping was consistently smaller and exhibited larger variations, which can be attributed to end effects at the non-cylindrical pore opening^20^.

**Figure 3.**
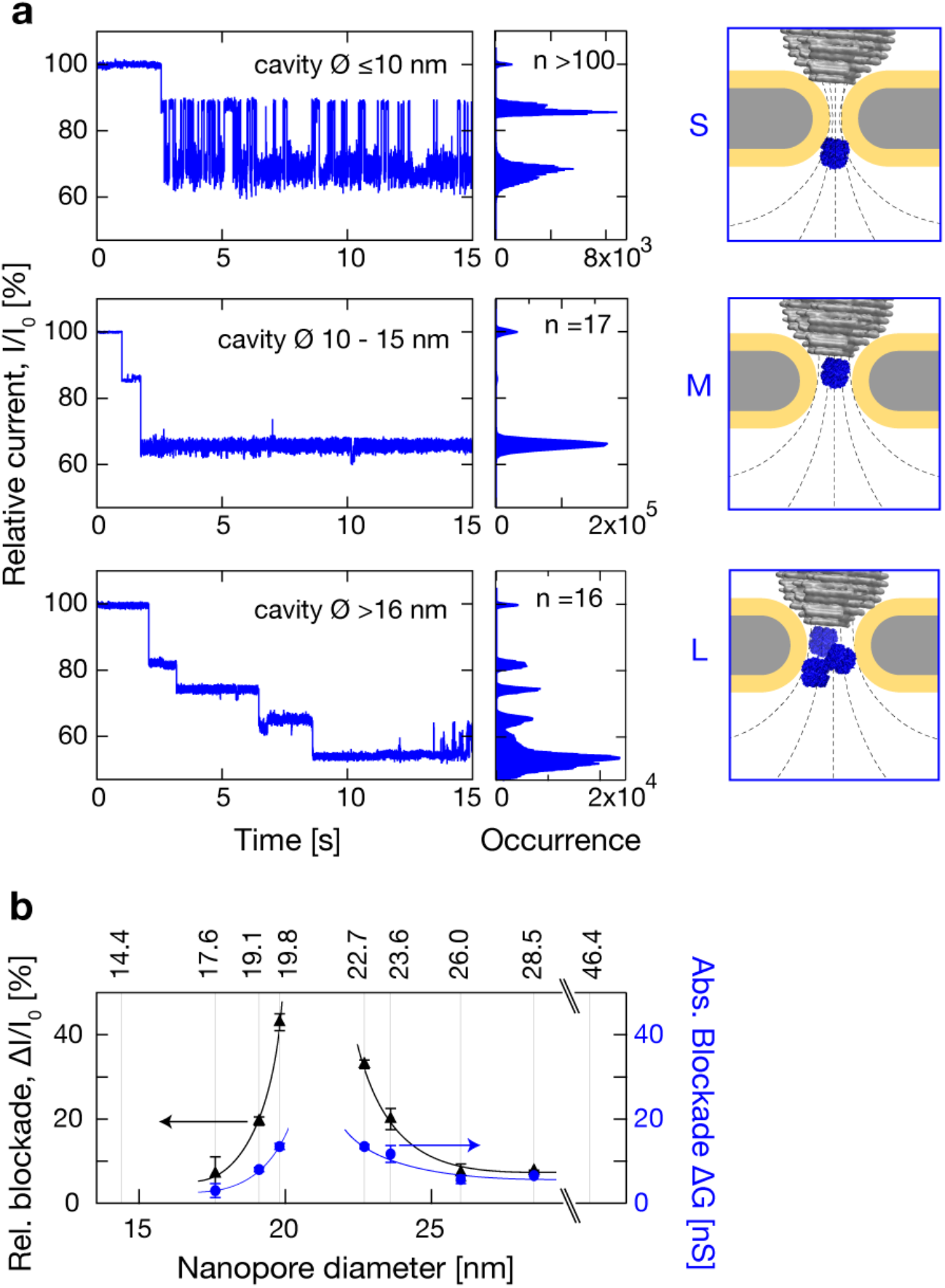
Pore size-dependence of NEOtrap signals. **a** Relative current traces and histograms showing ClpP trapping in three pore size regimes as illustrated on the right. From top to bottom: (S) a too small cavity leading to short trapping events where ClpP cannot enter the pore but merely blocks the entrance temporarily (here d_coated pore_= 9.5 nm); (M) a cavity fitting the protein size leading to long-term single-protein trapping (here d_coated pore_= 14.0 nm); (L) a cavity fitting multiple proteins leading to consecutive trappings of multiple proteins with discrete blockades (here d_coated pore_= 16.4 nm). Associated histograms combine many protein trapping events as specified. The protein-accessible cavity diameter is specified, i.e.: d_coated pore_ defined in the text. **b** Relative (black) and absolute (blue) current blockade levels of a single ClpP at 100mV, obtained over a large range of nanopore diameters: d_pore_= 14 to 46 nm (where d_pore_ is deduced from conductance before coating, see Supplementary Note 3.4). Error bars represent the full-width half maximum (FWHM) of Gaussian fits to current histograms over 10-30 origami docking events per datapoint. Lines are guides to the eye. The origami sphere in panel (a) was adapted from Ref.10.

The cavity dimensions accessible for protein trapping are thus defined by the size of the protein itself (at the small limit), and by the translocation of the 35 nm diameter origami sphere (at the large limit, Fig. 3b). As expected, the absolute conductance blockade per ClpP grew from the too-small-cavity regime (S) to the intermediate regime (M) which fully accommodates ClpP, thus causing the biggest blockade. However, the absolute conductance blockade dropped again from the intermediate to the large pore regime (L), from 11.7±2.0 nS at 23.6 nm pore diameter to 5.7±1.0 nS at 26 nm pore diameter. Microscopically, this may be explained by a sterically hindered ion flow when ClpP precisely fills the cavity, which causes an additional excluded volume for ions with associated water shells which adds to the total ClpP blockade.

The applied voltage affects the conductance blockade caused by the docked origami sphere differently compared to the one caused by the trapped protein, see Fig. 4a for data of two pore sizes and Fig 4b for corresponding current traces. The blockade levels of the trapped ClpP proteins stayed constant with applied voltage. This indicates that the protein stays intact, and protein unfolding by electrostatic or hydrodynamic forces can be excluded for the voltage range probed. In contrast, the current blockade levels of the docked origami sphere grew significantly larger at higher voltages. We attribute this to the DNA-origami sphere that is gradually pressed more tightly onto the nanopore with increasing voltage.

**Figure 4.**
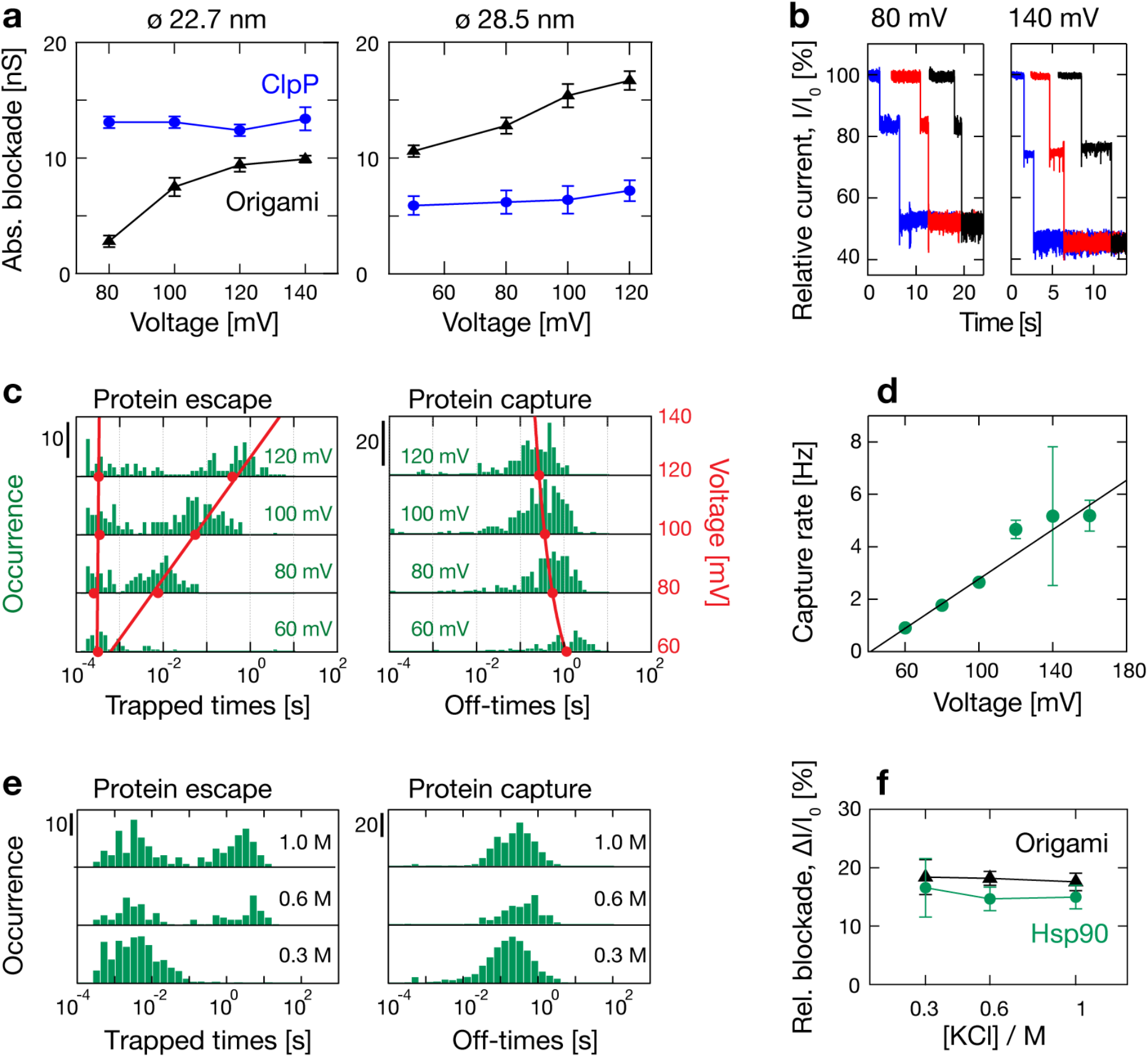
NEOtrap characteristics as a function of voltage and ionic strength. **a** Voltage dependence of ClpP (blue) and origami blockades (black) for two pore diameters as specified. Error bars represent the FWHM of Gaussian fits to current histograms of 10-30 docking and trapping events per datapoint. **b** Example current traces of the d_pore_ = 22.7 nm case in a, under the maximum and minimum voltage as specified. **c** Hsp90 protein trapping kinetics (from one origami docking) for a range of applied voltages, as specified. Scale bars specify number of counts. The protein trapped times (left) reproducibly split into two populations – highlighted with red lines, which are linear fits to the population centres (red dots) determined with lognormal fits – indicating two escape pathways (observed in ≥5 measurements per condition). The off-times (no protein trapped, right panel) form single-exponential distributions described by one voltage-dependent capture rate. The red line (and right axis) is a linear fit to the capture rates as a function of voltage. Corresponding dot plots are shown in Supplementary Fig. 4 and 5. **d** The protein capture rate grows linearly with applied voltage (here for multiple origami docking events). The black line is a linear fit with slope = 0.047±0.003 Hz/mV and y-axis intercept = -1.9±0.21 Hz. The reported rates and error bars were obtained as means and standard deviations from bootstrapping single-exponential fits to off-time histograms (see Methods). **e** Protein trapping kinetics for Hsp90 for different KCl concentrations of 0.3-1 Molar. Dwell-time histograms include multiple origami docking events. Scale bars specify number of counts. Similar to the data in panel c, the trapped times split into two populations and the off-times (no protein trapped) are single-exponentially distributed. Note that the capture rate barely changes with salt concentration. Long trapped events may be underrepresented by the recording time (here 20 s per origami docking, see current traces in Supplementary Fig. 6). Dwell-time distributions of minute-long recordings are shown in Supplementary Fig. 7. **f** No significant salt dependence was observed for the relative current blockades induced by the docked origami or trapped Hsp90. All Hsp90 data in this figure was measured in the presence of 5mM AMP-PNP.

Protein trapping kinetics provide further insight into the trapping mechanism of the NEOtrap. Here, we studied Hsp90 as a model protein, because the very low escape rate (<1/hour) of the stably trapped ClpP precluded sufficient event statistics. Figure 4c shows histograms of the protein trapped times (protein residence times) and the non-trapped off-times (empty cavity) for Hsp90, which inform on the escape and capture process, respectively. Two escape pathways are consistently observed, a fast escape process with a trapped time that is voltage independent, and a majority escape process that is slower, with a trapped time that grows exponentially with voltage. This exponential dependence is expected for a particle escape process that involves an energy barrier crossing as described by Boltzmann statistics, cf. Supplementary Note 3.3. The voltage-independent fast escape process may involve protein escape through the origami side as a result of temporal fluctuations in the docking of the sphere. Protein capture, on the other hand, exhibits a single-exponential off-time distribution with a linear voltage dependence of the capture rate (Fig. 4c right), as theoretically predicted. The linear fit of the capture rate in Fig. 4d, drops to zero at ∼40 mV, which is of the order of the minimal voltage required for permanent docking of the origami sphere (here ∼60mV). On the high voltage end, the stability of the lipid coating limits the experimentally accessible voltage range to ≤160 mV. A voltage of ∼100 mV was found to be optimal regarding signal-to-noise ratio and trapping kinetics in the pore diameter range from 20-25 nm.

Scanning three different ionic strengths (Fig. 4e), we find the longest trapped times under an intermediate to high KCl concentration (0.6 – 1 M), while only the short-trapped population is observed at 0.3M KCl (see also current traces in Supplementary Fig. 6). The capture rate, on the other hand, remains remarkably constant over the three ionic strengths. The relative blockade also remains unchanged within experimental uncertainty (Fig. 4f), which is further evidence for a non-invasive trapping mechanism that does not affect the protein stability, for neither ClpP nor Hsp90.

## Label-free detection of nucleotide-dependent conformational shifts in the chaperone Hsp90

Finally, we demonstrate the NEOtrap’s capacity for label-free detection of protein conformational heterogeneity and nucleotide-binding-induced conformational shifts in proteins, such as Hsp90 (Fig. 5). The diverse conformations of Hsp90 play a key role in the functional cycle of this molecular chaperone^21^, which is a central metabolic hub for protein homeostasis in the cell^22^ and an enabler of cancer adaption^23^. Figure 5a depicts various conformations of homo-dimeric Hsp90, its two N-terminal nucleotide binding sites, and the many non-covalent bonds that the nucleotide (here ATP) engages in, which critically stabilize multiple domains of the protein^24^. Figures 5b,c compare NEOtrap recordings of Hsp90 under four different nucleotide conditions: with 5 mM AMP-PNP, ATP, ADP, or in the absence of any nucleotide, termed ‘apo’. Clear differences are observed: in the presence of ATP or the non-hydrolysable ATP-analogue AMP-PNP, one dominant peak is observed at a low conductance blockade of -5.4 ± 0.2 nS or -5.3 ± 0.2 nS, respectively, while larger blockades are absent, which we interpret as Hsp90 prevailing in one compact conformation (in line with existing literature discussed below). Larger blockades were occasionally observed in the presence of ATP (see representative traces in Fig. 5b, and the shoulder in the ATP histogram in Fig. 5c). In the presence of ADP, more diverse blockade levels are detected, indicating a more diverse conformational ensemble populated by Hsp90. Lastly, in the absence of any nucleotide, where Hsp90 is missing essential stabilizing contacts in the N-domain, the blockade levels are distributed the most heterogeneously. Note also that the higher blockade levels exhibit larger fluctuations in the current traces (Fig. 5b), suggesting increased conformational dynamics in the millisecond range.

**Figure 5.**
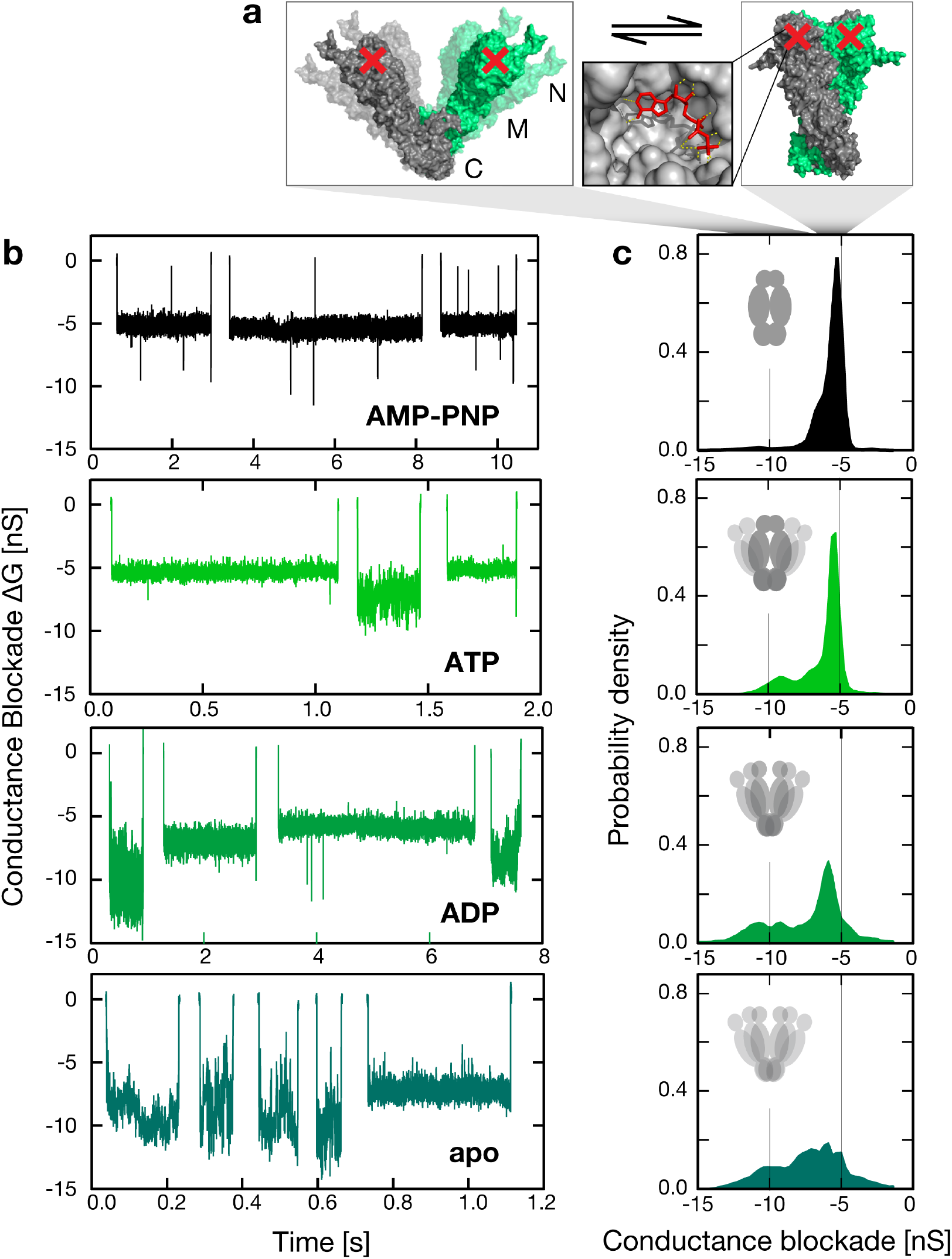
Label-free NEOtrap detection of nucleotide-dependent conformational shifts of the chaperone protein Hsp90. **a** Hsp90 in flexible open (left^26^) and compact closed conformations (right, pdb: 2cg9^24^) of the homo-dimeric chaperone protein Hsp90, consisting of an N-terminal nucleotide-binding domain (N), a middle domain (M), a C-terminal dimerization domain (C). Hsp90 alternates between these conformations depending on different nucleotide-binding states. The inset shows the many stabilizing contacts (yellow dashed lines) formed by the nucleotide in Hsp90’s Bergerat-type ATP-binding site. **b** Representative conductance blockade events caused by single label-free Hsp90 proteins under the specified nucleotide conditions: non-hydrolyzable AMP-PNP, ATP, ADP, apo. Hsp90’s conformational heterogeneity increases strongly from top (AMP-PNP, compact) to bottom (apo, flexible), in line with the reduced number of stabilizing nucleotide contacts, and with existing Hsp90 literature. **c** Histograms of 1000-4000 such current blockade events (integral normalized to unity) that show a strong nucleotide dependence, indicative of distinct protein conformations that are visualized using gray icons where dark represents frequent, and pale rare conformations, respectively. Scatterplots of conductance versus dwell time are provided in Supplementary Fig. 8. Less than 1% outlier events were excluded as described in Supplementary Fig. 9. The results were reproduced in three or more experiments using different measurement orders of the four nucleotide conditions.

The striking differences observed in these current recordings of unlabelled Hsp90 reveal different conformational states of the Hsp90 dimer: AMP-PNP binding strongly stabilizes Hsp90’s compact conformation^24–26^. Consistently, we observed one clear low-blockade peak. In the cases of ATP, ADP and apo, this structural rigidity and compaction is increasingly lost, and Hsp90 undergoes more and more excursions to multiple thermally accessible open conformations. Our results match with previous findings from smFRET^25,26^, electron-microscopy^27^, and biochemical results^28^. However, in contrast to these techniques, the NEOtrap measures proteins in a label-free manner, at room temperature, in solution, and at the single-molecule level. And unlike smFRET, the spatial resolution of the NEOtrap is not limited to a certain inter-dye distance (typically ≤9nm), and the observation time is not limited by photobleaching. The ability to distinguish nucleotide-dependent conformations in a protein complex demonstrates the great potential of the NEOtrap, opening the way for studying conformational dynamics in many protein systems.

## Conclusion

In conclusion, we presented the Nanopore Electro-Osmotic trap, which is capable of trapping single unmodified proteins irrespective of their net charge for up to hours while resolving their size and conformations with sub-millisecond time resolution. This represents a drastic increase in single-protein observation time by a factor of >10^6^ compared to previous solid-state nanopore studies^19,29^. A DNA-origami sphere was used to induce local electro-osmotic flows at will, which creates the trapping potential for the NEOtrap. Voltage inversion can be applied as a clean sweep to trap a new protein molecule in the lipid-passivated nanocavity whose size can be freely chosen between a few nm and 30 nm, i.e., in a size range very relevant for proteins and other molecules, nanoparticles, quantum dots, etc. The NEOtrap features a linear mass dependence of the current blockade for globular proteins of 54kDa to 340kDa; can discriminate between different orientations for disk-shaped ClpX; and, most remarkably, distinguishes nucleotide-dependent conformations of the chaperone protein Hsp90. The NEOtrap offers a widely applicable electrical sensing strategy that addresses a central shortcoming of nanopore detection, viz., the impractical high-speed translocation that prohibits single-protein characterization. As a result, the NEOtrap has the potential to become a label-free alternative to classical single-molecule techniques like fluorescence and force spectroscopies. Due to its technical simplicity and multifaceted applicability, we anticipate that the NEOtrap will find wide-spread application in the study of diverse protein systems, and thus evolve into a new tool in the single-molecule toolbox for studying protein dynamics.

## METHODS

Glass chips with free-standing 20nm silicon nitride membranes were purchased from Goeppert LLC (Philadelphia, USA). Nanopores were drilled by TEM as previously described^8^. The chips were rinsed with MilliQ water, ethanol, acetone, isopropanol, and plasma cleaned (SPI Supplies, West Chester, USA) before the assembly in a custom-made PEEK flowcell with cis and trans buffer compartments, placed in a Faraday cage, and connected to an Axopatch 200B amplifier and Digidata 1550B digitizer (both Molecular Devices, San Jose, USA) using Ag/AgCl electrodes (silver wire chloridized in household bleach). All pores were wetted with MilliQ water, before measuring the conductance in 1M-KHM buffer (1M KCl, 50mM Hepes, 5mM MgCl_2_, pH7.5) to obtain a measured nanopore diameter as described in Supplementary Notes 3.4.

The nanopore passivation procedure was adapted from Ref.19: POPC (Avanti Lipids, Alabaster, USA) in chloroform was aliquoted in glass vials, dried for 2-4 hours in vacuo, either stored at -20°C (≤ 1month), or resuspended to 1mg/ml in 600KHM buffer (600mM KCl, 50mM Hepes, 5mM MgCl_2_, pH7.5), and vortexed for 10min. 50µl lipid suspension were applied to the cis compartment of a chip equilibrated in 600KHM, while applying a continuous zig-zag potential (peaks at ±50mV, 5Hz repetition rate). This step was repeated after 5 min. (If no conductance change was observed, 1M-KHM was added to the trans compartment to osmotically drive lipid vesicles to the pore.) After a total of 10-15min incubation time, the flowcell was disconnected and immersed in 0.5l MilliQ water for 10-15min. Next, the (externally dried) flowcell was reconnected to the amplifier and the nanopore flushed with 200µl MilliQ water in cis and trans, followed by the desired measurement buffer. A reproducible 1-step conductance drop indicated a stable and consistent coating. About 10% of the pores could not be stably coated in this way and were not used further. Other chips were re-coated and re-used several times using this procedure.

Nanopore experiments were controlled using ClampEX (Molecular Devices), and data analysis was performed using self-written code in Igor Pro v6.37 (Wavemetrics, Portland, USA). The open-source NeuroMatic binary file importer was used^30^. Median-aware decimation was used to exploit oversampling: e.g., data recorded at 500kHz sampling was decimated to 5kHz by replacing each 100 samples by their median value. Threshold criteria were used for event detection. Bootstrapping was used to estimate uncertainties of (i) kinetic rate constants from single-exponential fits and (ii) histogram peak positions as specified: for a dataset of size n, 10n subsets of size n were randomly chosen with replacement, individually evaluated, and the mean and standard deviations across all subsets are reported. Protein structures were visualized with the PyMOL Molecular Graphics System, Version 2.0.6, Schrödinger, LLC (New York, USA).

The DNA-origami sphere was described previously^10^. Details were kindly provided by tilibit nanosystems GmbH (Munich, Germany). Folding reaction mixtures contained 7560 nt-long scaffold DNA at 50 nM final concentration and 227 staple strands at 175 nM final concentration, for each strand. The folding buffer contained 5 mM TRIS, 1 mM EDTA, 5 mM NaCl (pH 8) and 20 mM MgCl_2_. Folding reaction mixtures were subjected to thermal annealing ramps using TETRAD (MJ Research, now Biorad, Hercules, USA) thermal cycling devices: 15 minutes at 65°C, followed by one-hour intervals for each temperature, starting at 64°C down to 48°C, decreasing by 1°C every step. The folding reaction mixtures were then incubated at 20°C before purification steps. Excess staple strands were removed by ultrafiltration (Amicon Ultra 0.5 ml Ultracel filters, 50K) with buffer containing 5 mM TRIS, 1 mM EDTA, 5 mM NaCl and 5 mM MgCl_2_ ^31^: Add 0.5 ml of buffer and centrifuge at 10k G for 3 minutes at 25°C. Discard flow through. Add 0.1 – 0.2 ml of folded object sample and 0.3 – 0.4 ml of buffer and centrifuge 10k G for 5 minutes at 25°C. Discard flow through. Repeat step 2 for 3 more times. Remove filter inset, place upside-down into a new tube and centrifuge at 10k G for 3 minutes at 25°C as a sample retrieving step.

ClpP and ClpX were expressed and purified in-house as previously described^32^. Hsp90 was a kind gift of Bianca Hermann and Thorsten Hugel^25^. Avidin (mono-valent SAe1D3^33^) was a kind gift of Mark Howarth.

Unless stated differently all measurements were performed under 100 mV bias, 500kHz sampling, 100kHz low-pass filter (4-pole internal Bessel filter), in 600KHM, at room temperature of 21±1°C. If not stated differently chemicals and nucleotides were purchased from Sigma Aldrich (Zwijndrecht, The Netherlands).

## ACKNOWLEDGEMENTS

Hsp90 was a kind gift of Bianca Hermann and Thorsten Hugel. Avidin was a kind gift of Mark Howarth. ClpP and ClpX plasmids were a kind gift of Chirlmin Joo. We thank Xin Shi and Alessio Fragasso for discussions, Eli van der Sluis for discussions and protein purification, Meng-Yue Wu and Frans Tichelaar for TEM drilling. The work was funded by NWO-I680 (SMPS) and supported by the NWO/OCW Gravitation program NanoFront and the ERC Advanced Grant 883684. SS acknowledges the Postdoc.Mobility fellowship no. P400PB_180889 by the Swiss National Science Foundation. This work was supported by a European Research Council Consolidator Grant to H.D. (GA no. 724261), the Deutsche Forschungsgemeinschaft through grants provided within the Gottfried-Wilhelm-Leibniz Program (to H.D.).

## AUTHOR CONTRIBUTIONS

S.S. and C.D. conceived the project. S.S. performed all nanopore experiments, analyzed the data, and purified proteins. H.D. and P.S. advised on DNA origami, and P.S. folded and characterized it. S.S. wrote the manuscript with C.D. All authors discussed the results and commented on the manuscript.

## ADDITIONAL INFORMATION

**Supplementary information** is available in the online version of the paper.

**Reprints and permissions information** is available online at www.nature.com/reprints.

**Correspondence and requests for materials** should be addressed to C.D.

