## Supplementary Information for "Nanopore electro-osmotic trap for the label-free study of single proteins and their conformations"

**4 Supplementary References** **14**

### 1 Supplementary Figures

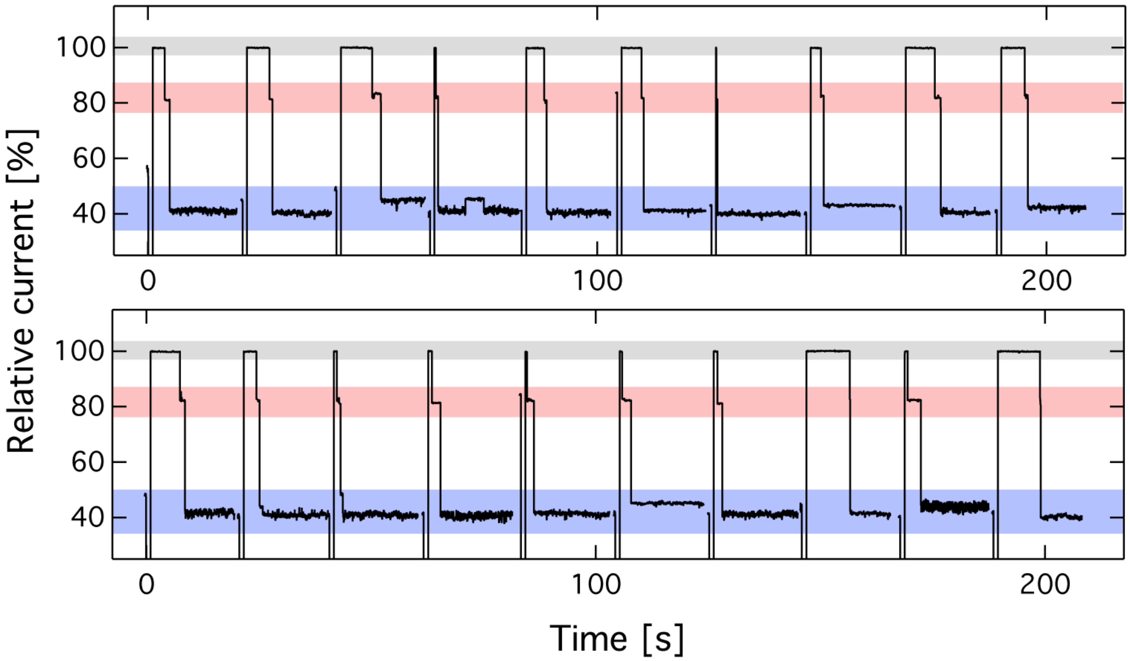

**Supplementary Figure 1**: Current recordings of ClpP protein trapping events. Color code as in Fig. 1 for the open-pore current, the origami-docking level and the protein-trapped current level (top to bottom). The data was recorded in series at 100mV, with voltage inversion (-50mV) every 20s to release the origami and renew the trap; using 100pM origami spheres, 10nM ClpP, in 600mM KCl, 50mM Hepes, 5mM $MgCl_2$, pH7.5, using a nanopore diameter of 20nm.

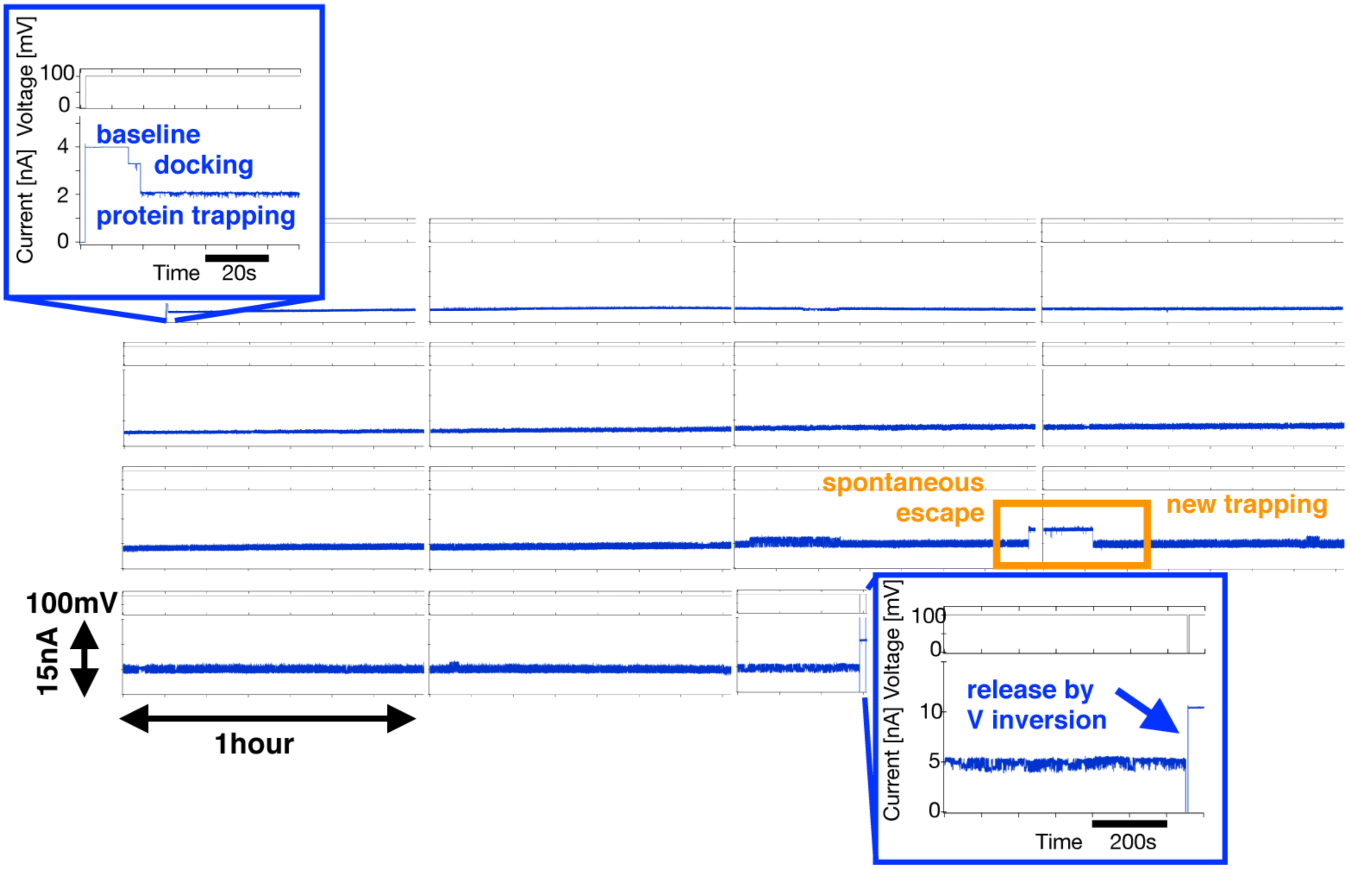

**Supplementary Figure 2**: Longterm trapping of ClpP. Non-corrected current recording with zoom view showing the usual pattern: open-pore baseline, followed by origami docking, and ClpP trapping. A spontaneous ClpP escape happens after 11h, followed by a new trapping as specified in the orange box. After another trapping for >3h ClpP is released manually by voltage inversion to -100mV, which recovers the low-noise baseline. Data was measured with 100pM origami spheres, 10nM ClpP, in 600mM KCl, 50mM Hepes, 5mM MgCl2, pH7.5, using a nanopore diameter of 21nm, and 1kHz filter. Two cut eppendorf tubes filled with water and inverted to cover both buffer reservoirs were used to improve longterm stability. Still the conductance and noise increased over the long time range of hours, likely owing to the limited pore coating stability, buffer conductivity increase by residual evaporation, and electro-chemical effects at the Ag/AgCl electrodes.

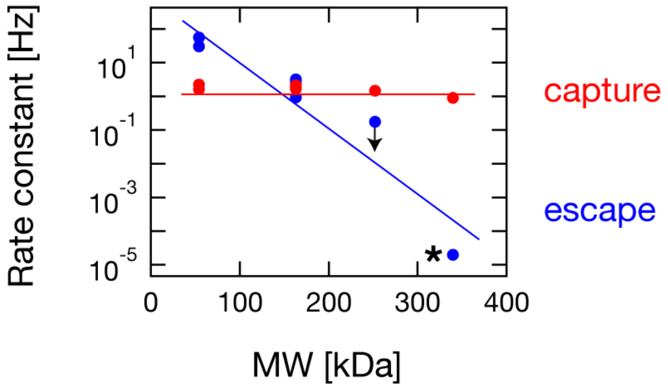

**Supplementary Figure 3**: Protein capture and escape rate constants as a function of molecular weight, as obtained from single-exponential fits to cumulative dwell-time histograms. The lines are guides to the eye, indicating the expected constant trend of the capture rates and exponential dependence of the escape rates. The black arrow indicates the overestimated ClpX escape rate due to the finite minute-long recording time. The black asterisk specifies an estimate of the ClpP escape rate based on the rare spontaneous escape shown in Fig. 1d. Data was obtained using 100pM origami spheres, 10nM respective protein, in 600mM KCl, 50mM Hepes, 5mM $MgCl_2$, pH7.5, using a nanopore diameter of 20nm.

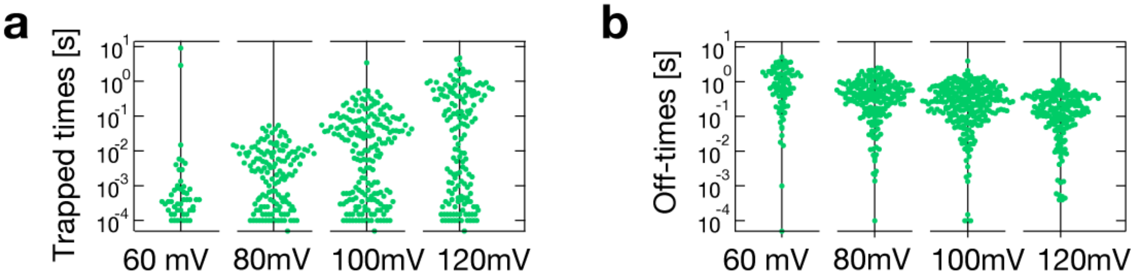

**Supplementary Figure 4**: Representative voltage-dependence of the Hsp90 protein's trapped times (a) and the un-trapped 'off-times' (b), for one single origami sphere docked onto the nanopore (shown in Fig. 4c). The point-cloud's total x-extension is a measure of occurrence. The data was measured with 100pM origami spheres, 10nM Hsp90, 5mM AMP-PNP, in 600mM KCl, 50mM Hepes, 5mM $MgCl_2$, pH7.5, using a nanopore diameter of 23nm.

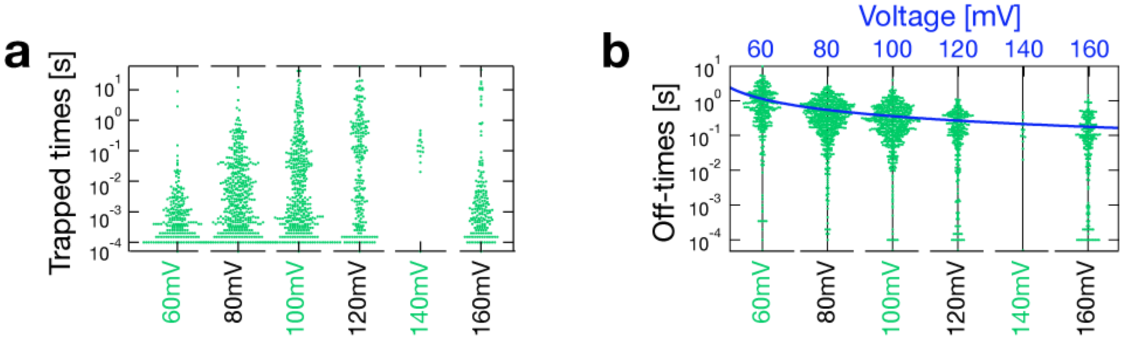

**Supplementary Figure 5**: Voltage-dependence of the Hsp90 protein's trapped times (a) and the un-trapped 'off-times' (b), obtained from multiple origami docking events, displayed as point clouds. The blue solid line in (b) is the reciprocal of the linear fit to the capture rate constants ($k = 1/\tau$) as a function of voltage, as shown in Fig. 4d. The point-cloud's total x-extension is a measure of occurrence. The data was measured with 100pM origami spheres, 10nM Hsp90, 5mM AMP-PNP, in 600mM KCl, 50mM Hepes, 5mM $MgCl_2$, pH7.5, using a nanopore diameter of 23nm.

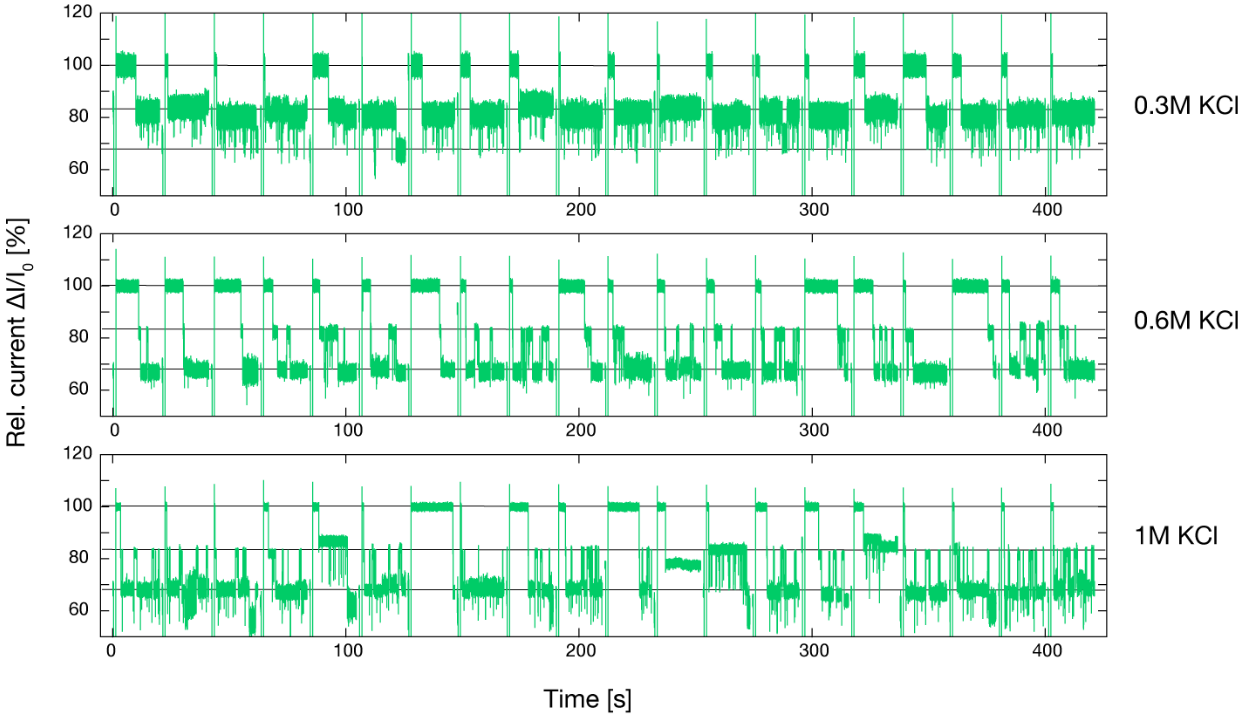

**Supplementary Figure 6**: Current recordings of twenty origami docking events with subsequent Hsp90 protein trappings for three different salt concentrations as specified. Three horizontal lines indicate (from top to bottom) the open-pore current, the origami-docking level and the protein-trapped current level. Data was recorded in series at 100mV, with voltage inversion (-50mV) every 20s to release the origami and renew the trap. The data was measured with 100pM origami spheres, 10nM Hsp90, 5mM AMP-PNP, in 600mM KCl, 50mM Hepes, 5mM $MgCl_2$, pH7.5, using a nanopore diameter of 21nm.

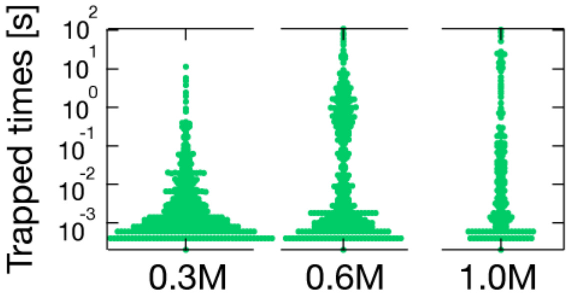

**Supplementary Figure 7**: Salt dependence of the protein trapped times obtained from multiple origami dockings in 2-10 minutes long recordings. Note that the recording time limits the observed long events at maximum 100s. The point-cloud's total x-extension is a measure of occurrence. The data was measured with 100pM origami spheres, 10nM Hsp90, 5mM AMP-PNP, in the indicated KCl concentration plus 50mM Hepes, 5mM $MgCl_2$, pH7.5, using a nanopore diameter of 23nm.

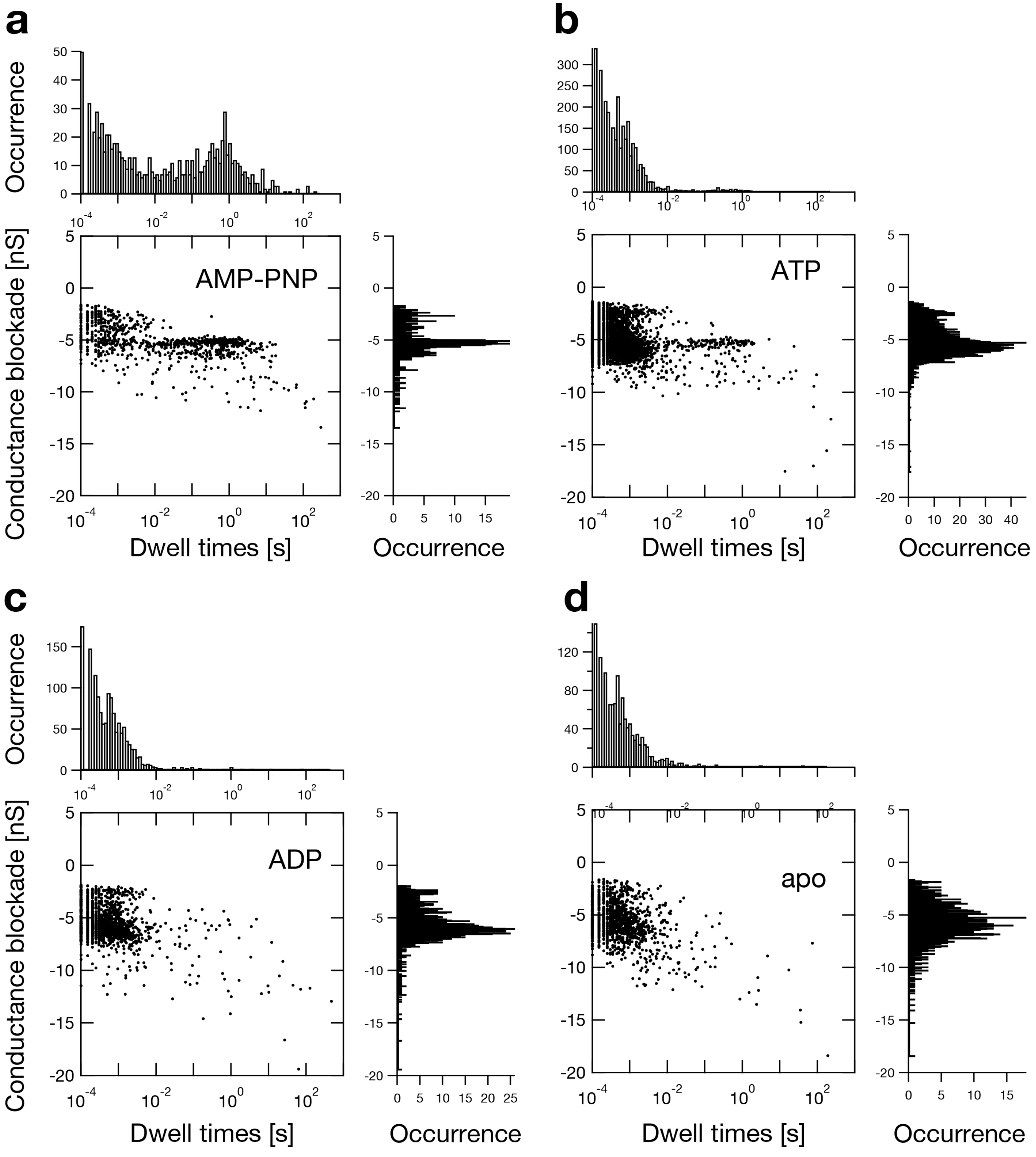

**Supplementary Figure 8**: Scatter plots of conductance blockade vs. dwell time, for Hsp90 under four nucleotide conditions (specified in a,b,c,d, respectively) along with 1D histograms for both dimensions. The data was measured with 100pM origami spheres, 10nM Hsp90, 5mM of the indicated nucleotide, in 600mM KCl, 50mM Hepes, 5mM $MgCl_2$, pH7.5, using a nanopore diameter of 24.5nm.

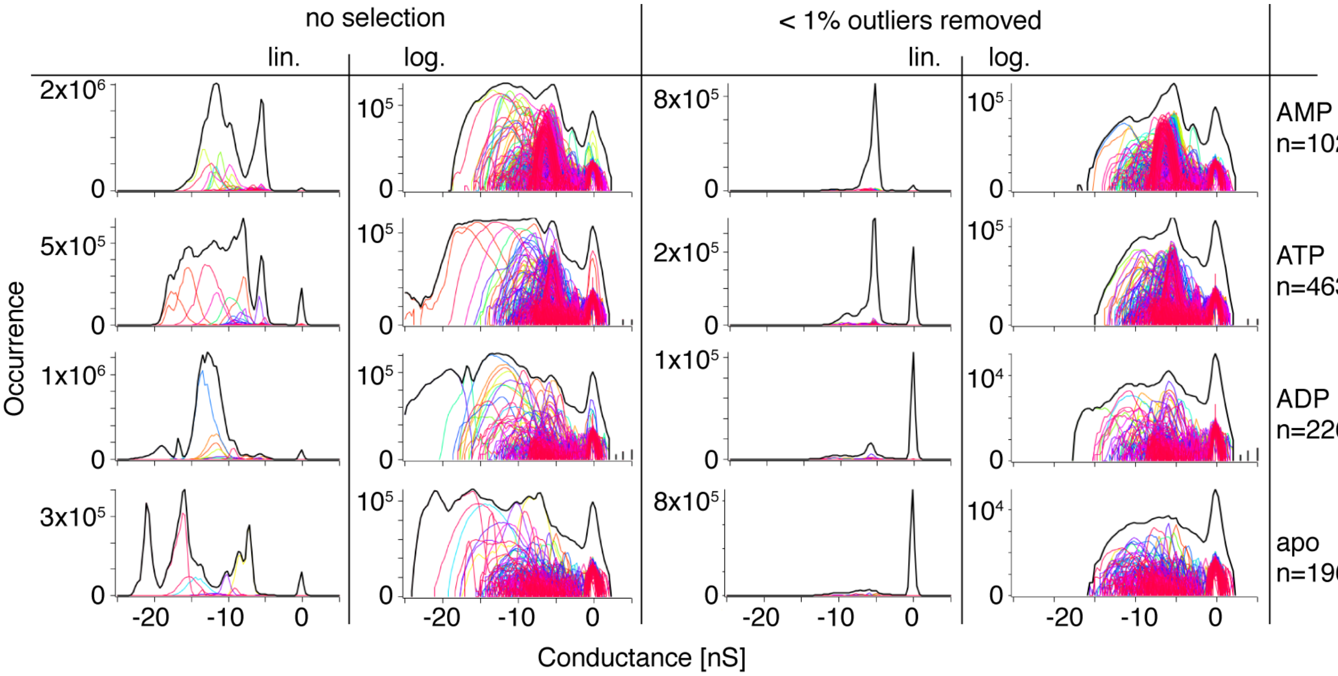

**Supplementary Figure 9**: Report of the event selection performed on the Hsp90 data in Figure 5: less than 1% outlier events were excluded. Outliers were identified based on 9899 single-event histograms, shown here in different colours along with the total histogram for each condition in black. About five to ten individual events caused individual peaks unrelated to the ensemble as seen on the left in linear (lin.) and logarithmic (log.) scale. The final histograms (right hand side) were obtained from the remaining 99% of events.

### 2 Supplementary Table

**Supplementary Table 1**: Mass, isoelectric point (pI), and charge (at pH7) of the proteins used in this work. All values represent sequence-based calculated numbers including usual purification tags (e.g. 6His), as reported in Geneious Prime 2020.0.5. yHsp90-wt was used in Figure 2 only.

| Protein | mass [Da] | pI | charge [$q_e$] |
|---|---|---|---|
| ClpP (14-mer) | 339'542 | 6.21 | -55.30 |
| ClpX (linked hexamer) | 252'262 | 4.61 | -100.87 |
| yHsp90-wt (dimer) | 162'812 | 4.53 | -75.14 |
| yHsp90-zip (dimer) | 174'962 | 4.67 | -66.92 |
| Avidin (tetramer) | 54'293 | 5.32 | -12.55 |

### 3 Supplementary Notes

The cartoon of the NEOtrap in Supplementary Figure **10** specifies basic terms used below for a theoretical description of the electro-osmotic flows, and the resulting trapping potential defining the capture and escape rates.

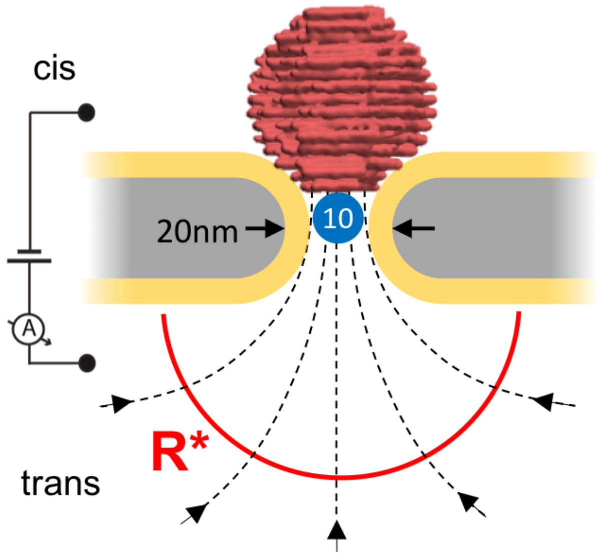

**Supplementary Figure 10**: The NEOtrap forms upon docking of an origami sphere (red, 35nm diameter) onto a lipid-coated (yellow) silicon nitride (gray) nanopore by positive forward voltage (cis to trans, as specified). The origami acts as an negatively charged sponge, which is ion-permeable and therefore highly electro-osmotically active: driven by the electric field, positive counter-ions flow through the origami from trans to cis, causing a stationary water flow in the same direction. This hydrodynamic velocity field (black dashed lines) creates a trapping potential. The capture radius R* (red bow) defines the half sphere within which particles (blue, e.g. 10nm diameter) get trapped. Once trapped, a particle can be observed up to hours - depending on size, voltage, and ionic strength.

#### 3.1 Electro-osmotic velocity

The electro-osmotic velocity $v_{eo}$ is a function of the zeta potential $\zeta$ of the charged object (here the origami sphere), the applied voltage causing the electric field $E$, the viscosity $\eta$, and $\epsilon$ the electric permittivity [1, 2]. We consider stationary conditions, i.e. a constant voltage, ion concentration, etc.

$$v_{eo}^{\circ} = \frac{\epsilon E \zeta}{\eta} \tag{1}$$

The $\circ$ symbol is a reminder that Equation (1) gives the basic electro-osmotic flow velocity - an overestimation for the conditions considered here: in particular, it neglects that the origami sphere is a nano-porous medium with incomplete charge screening internally, and furthermore it neglects any friction at the lipid-coated pore wall, and finally it ignores the local decay of the electric field over the size scale of the sphere. Equation 1 yields an upper limit for the flow velocity of $v_{eo}^{\circ} = 50$ mm/s for: $E = 100mV/30nm = 3.\overline{3} \cdot 10^6$ V/m, $\epsilon_r = 73$ [3], a measured $\zeta = -23.4$ mV, a measured $\eta = 0.96$ cP $= 9.6 \cdot 10^{-4}$ kg/s/m, all in 600mM KCl, 5mM $MgCl_2$, 50mM Hepes, pH7.5 and at room temperature of $21 \pm 1°$C. The origami zeta potential and viscosity were measured by dynamic light scattering on a Zetasizer Nano ZS (Malvern, Kassel D), according to the suppliers protocol.

#### 3.2 The capture radius and capture rate

The maximum solvent velocity *inside* the nanopore, $v_{eo}^{\text{in}}$, with diameter $a$ creates a flux $J_0$, and - by continuity - decreasing velocities $v_{eo}(r)$ at increasing distances $r$ away from the nanopore [4]:

$$J_0 = \int_0^a v_{eo}^{\text{in}}\, 2\pi r dr = \pi\, v_{eo}^{\text{in}}\, a^2 \tag{2}$$

$$v_{eo}(r) = \frac{J_0}{2\pi r^2} = \frac{a^2}{2\, r^2}\, v_{eo}^{\text{in}} \tag{3}$$

Next, from the diffusion coefficient D, we know the average time $t$ it takes for a particle with radius $r_{\text{part.}}$ to travel a distance $x$ by diffusion. Following the ansatz for electro-phoretic capture by Wanunu *et al.* [5], we define a 'diffusive velocity' $v_{\text{diff}} = r/t$ in Equation 6.

$$D = \frac{\langle x^2 \rangle}{2t} = \frac{k_B T}{6\pi\eta r_{\text{part.}}} \tag{4}$$

$$t = \frac{\langle x^2 \rangle}{2D} \approx \frac{r^2}{2D} \tag{5}$$

$$v_{\text{diff}} = \frac{r}{t} = \frac{2D}{r} \tag{6}$$

The capture radius $R^*$ is then defined as the distance away from the nanopore where the electro-osmotic velocity (Equation (3)) and the mean diffusive velocity (Equation (6)) are equal. That is, it represents the border between the diffusion-dominated and drift-dominated regimes:

$$v_{\text{diff}}(r) = v_{eo}(r) \qquad | \, r = R^* \tag{7}$$

$$\frac{2D}{R^*} = \frac{a^2}{2\,(R^*)^2}\, v_{eo}^{\text{in}} \tag{8}$$

$$R^* = \frac{a^2}{4D}\, v_{eo}^{\text{in}} \tag{9}$$

This capture radius defines a half-sphere around the nanopore opening (at the 'trans' side in Suppl. Fig. **10**), inside which particles get captured by the trapping potential. The capture rate, can then be defined as the rate at which particles enter the half sphere defined by the capture radius. With the particle concentration $c$, the capture rate constant in units of $\left[\frac{\text{mol}}{s}\right]$ becomes:

$$k_{on} = 2\pi R^* D c = \frac{\pi}{2} a^2 c\, v_{eo}^{\text{in}} = \frac{\pi}{2} a^2 c \left(\frac{\epsilon E \zeta}{\eta}\right) \tag{10}$$

where in the last equation, we assumed $v_{eo}^{\text{in}} = v_{eo}^{\circ}$, i.e. the maximum flow velocity.
The capture rate thus scales:

- linearly with applied voltage and particle concentration $c$
- quadratically with the pore radius $a$
- inversely with viscosity $\eta$.

It is worth noting that the capture rate does *not* depend on the particle radius (or the molecular weight), although the capture radius does: smaller and thus faster diffusing particles have a smaller capture radius, but they reach it more frequently than larger slower diffusing particles, thus cancelling the particle size dependence of the capture rate.

#### 3.3 The trapping potential and escape rate

The potential energy well of the NEOtrap is formed by the Stokes drag due to the electro-osmotic flow field:

$$\vec{F}_{\text{stokes}} = 6\pi r_{\text{part.}} \eta \, \vec{v}_{eo} = \frac{k_B T}{D} \vec{v}_{eo} \tag{11}$$

where the last equation makes use of the Stokes-Einstein equation.

The depth of the potential energy minimum is given by the work required to pull a particle out of the potential well ($r = 0$) to the capture radius $R^*$. For convenience, we (i) set $r = 0$ at the center of mass of the particle, (ii) we choose to pull along the z-axis neglecting variation in x and y, and (iii) we sum up the 1D integration inside and outside the nanopore:

$$E_{\text{trap}} = \int_{r=0}^{R^*} F_{\text{stokes}} dr = \int_{r=0}^{l} F_{\text{stokes}}^{\text{inside}} dr + \int_{l}^{R^*} F_{\text{stokes}}^{\text{outside}} dr \tag{12}$$

Inside the nanopore (where the electric field lines are, to first approximation, parallel) the flow velocity is constant along the z-axis:

$$E_{\text{trap}}^{\text{inside}} = \int_{r=0}^{l} F_{\text{stokes}}^{\text{inside}} dr = \frac{k_B T}{D} v_{eo}^{\text{in}} \, \ell \tag{13}$$

Outside the nanopore the velocity drops as a function of the pore distance $r$. We use the right most expression of Equation (11) together with Equation (3): $v_{eo}(r) = a^2/(2r^2) \cdot v_{eo}^{\text{in}}$:

$$E_{\text{trap}}^{\text{outside}} = \frac{k_B T}{D} \int_{\ell}^{R^*} v_{eo}(r) \, dr = k_B T \left( \frac{a^2 \, v_{eo}^{\text{in}}}{2D} \right) \int_{\ell}^{R^*} \frac{1}{r^2} \, dr = k_B T \left( \frac{a^2 \, v_{eo}^{\text{in}}}{2D} \right) \left( \frac{1}{\ell} - \frac{1}{R^*} \right) \tag{14}$$

This yields the total trapping potential:

$$E_{trap} = k_B T \, \frac{v_{eo}^{\text{in}}}{2D} \left( \frac{a^2}{\ell} - \frac{a^2}{R^*} + 2\ell \right) = 2 k_B T \, R^* \left( \frac{1}{\ell} - \frac{1}{R^*} + \frac{2\ell}{a^2} \right) \tag{15}$$

To leave the NEOtrap, a particle has to cross a potential barrier with $E = E_{trap}$, and the escape rate constant follows Boltzmann statistics:

$$k_{\text{off}} = A \cdot \exp\left( -\frac{E_{\text{trap}}}{k_B T} \right) = A \cdot \exp\left[ \frac{v_{eo}^{\text{in}}}{2D} \left( \frac{a^2}{R^*} - \frac{a^2}{\ell} - 2\ell \right) \right] = A \cdot \exp\left[ 2R^* \left( \frac{1}{R^*} - \frac{1}{\ell} - \frac{2\ell}{a^2} \right) \right] \tag{16}$$

where $A$ is an Arrhenius-type frequency factor.

The escape rate constant thus scales:

- exponentially with voltage ($R^* \propto V$)
- with the nanopore radius as $\exp(-a^2)$ since $R^* \propto a^2$
- with the particle radius as $\exp(-r_{part.})$ since $R^* \propto 1/D \propto r_{part.}$

##### 3.4 Conductance-derived nanopore diameter

The nanopore diameter was determined from the conductance at 1M KCl using the following formula, considering surface charge [6] and access resistance [7]:

$$\frac{1}{G_0} = R_{tot} = \frac{L}{\pi D} \frac{1}{\sigma \cdot D/4 + n_{\text{surf}} \cdot \mu_{K^+}} + \frac{1}{\sigma D} \tag{17}$$

with $L$ the empirical nanopore length previously determined to 8.6 nm for TEM drilled pores in 20 nm silicon nitride membranes [7]; $D$ the nanopore diameter; $\sigma = 10.5\ S/m$ the bulk conductivity of 1M potassium chloride [8]; $n_{\text{surf}} = 0.02\ C/m^2$ the surface charge density of silicon nitride at pH7-8 [9]; $\mu_{K^+} = 7.6 \cdot 10^{-8}\ m^2/(Vs)$ the ion mobility of potassium [10]. The formula was validated with TEM images.
